# Fitting multiple bacterial growth data using continued fraction of straight lines

**DOI:** 10.1101/2025.01.27.634991

**Authors:** I S Shruti, S Vijay Prakash

## Abstract

The growth of a population is the net result of growth and decline in the number of individuals over time. A population grows when the increase in number of individuals is more than the decrease and declines in the opposite scenario. In other words, the growth rate of a population is influenced by two opposing factors, a growth promoting factor and a growth restricting factor. In this work, we estimate growth rates by applying a biological growth model that is based on the continued fraction of straight lines with two parameters *a* and *m*. The parameters *a* and *m* represent nonlinear i.e. growth restricting and linear i.e. growth promoting parts of the model, respectively. To fit this model, we use a publicly available dataset that exhibits the growth of three different strains of bacteria depending on the concentration gradient of the antibiotic Tetracycline. We also propose a method to automatically estimate growth rates for large-scale applications. Finally, a growth coordinate system with *a* and *m* as the axes is used to interpret the estimations.

**Importance:** In this work, multiple bacterial growth data has been fitted with a model based on continued fraction of linear growth. The importance of this work lies in the fitting of both growth and death phase with a single model. Rather than modeling growth with differential equations, this model uses algebraic expressions. Therefore, the growth rates are obtained directly from these expressions after fitting. Several of these models can be superposed, and more flexible fits can be obtained based on requirements. There are two parameters that play key roles in fitting. Their values can be expressed on planar plots which are useful to compare multiple growth data. Thus, this methodology provides simpler, generic, flexible and more interpretable growth models.

## 1 Introduction

Generally, growth curves are modeled by theoretically relating instantaneous growth rate with the instantaneous number of individuals using differential equations [1, 2, 3] resulting in multi-parametric logistic models. Another approach is to utilize empirical models such as polynomials [4, 5] to model growth curves. In recent years, deep learning has also been used to estimate growth rates, particularly for bacterial growth [6].

Growth curves can also be modeled as a continued fraction of linear growth in time [7]. This way of modeling growth has only two parameters *a* and *m*, out of which *m* theoretically represents the linear growth rate, while the other parameter *a* is tuned to fit the model to the growth data. So, effectively, a one-parametric model is introduced with the continued fraction approach, henceforth referred to as *a* − *m* model. According to this model [7]:

1. Biological growth is linear but restricted (or regulated) in a nonlinear manner.
2. The nonlinear variation of growth is symmetric about the point of maximum growth.
3. At the point of maximum growth, restrictions are negligible. However, the restrictions become more pronounced as we move away from the point of maximum growth rate.

### 1.1 Bacterial growth

The environment and its own physiology greatly influence the division of a bacterium [6]. The multiplication rate of bacteria is important in pathological studies, food safety and veterinary sciences [8, 9, 10]. Variations in the growth rate of a bacterial culture give rise to successive phases of growth [11]. Considering the overall growth of the culture, there are mainly two phases. One where the growth rate is either zero or positive and the other is the phase of decline where the growth rate is negative. In this work, we consider a bacterial growth data set and fit the *a* − *m* model to interpret the data.

#### About the dataset

The publicly available data set used in this work is given by Claudia Seiler from TU Dresden as part of a series of plate reader experiments [12]. Growth of three strains of bacteria, depending on the concentration of the antibiotic Tetracycline, was studied with values obtained for two replicates of the experiment. The data set consists of five columns. First one identifies the strain of bacteria namely, Donor (D), recipient(R) and Transconjugate (T). The next column gives whether it is the first or second replicate of the trial. The third column stands for the concentration of Tetracycline. The fourth and fifth column represent time in hours and bacterial population measured as optical density, respectively.

## 2 The *a* − *m* growth model

The *a* − *m* model is given by [7]

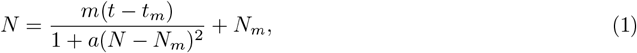

where *N*_*m*_ is the population size at time *t*_*m*_ of maximum growth rate *m*, with *N* as the actual population size at time *t*. Here, the term *a*(*N* − *N*_*m*_)^2^ represents the nonlinear restrictions to growth through the positive parameter *a*. The above equation can also be written as

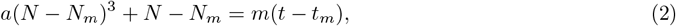

which provides us the rate of growth given by

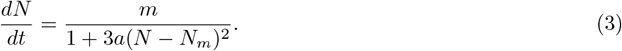

The growth rate is maximum at *N* = *N*_*m*_. Thus, *m* is the maximum growth rate of the model and *a* regulates this growth rate. The real solution of Eqn. (1) with *a >* 0 is given by

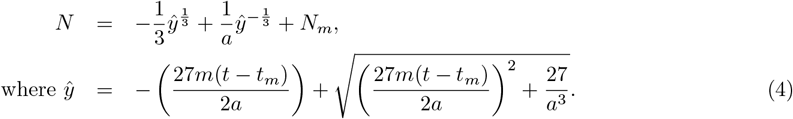

Eqn. (2) can also be expressed as a continued fraction [13]

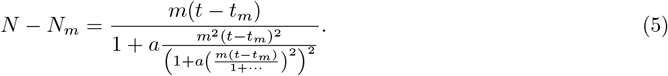

Eqns. (4) and (5) represent the same *N*. Thus, the continued fraction of linear growth *N* − *N*_*m*_ = *m*(*t* − *t*_*m*_) leads to the solution given by Eqn. (4).

The three forms of *a* − *m* model are useful in several ways as summarized below:

- 1^st^ form: Eqn. (2) provides us with convenient derivative expressions (Eqn. 3).
- 2^nd^ form: Eqn. (4) being the analytical solution helps us to numerically estimate the parameters in a straight-forward manner.
- The 3^rd^ continued fraction form (Eqn. 5) helps us to interpret the model. The interpretation is that population size grows linearly with time but restricted (or regulated) at various nonlinear levels. This results in infinitely regulated linear growth.

### 2.1 Fitting procedure

The fitting procedure involves choosing the origin i.e., (*t*_*m*_, *N*_*m*_) of Eqn. (4) and fitting the *a* − *m* model using the lmfit python package [14]. There are two ways of choosing the origin. One way is to manually select data points that form an S-shape within the entire curve and fit the model in such a way that the Bayesian information criterion is minimal[7]. And then we take the mean of selected points to be the origin (*t*_*m*_, *N*_*m*_) as shown in Fig. (2a). Although this gives more precise estimations of *a* and *m*, the decline phase has to be excluded to obtain them. The second way is to find the highest absolute slope in the growth data so that it includes the change due to growth or decline. The slope at *i*^*th*^ data point is given by

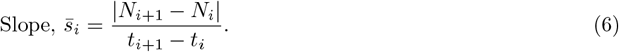

Then we select the pair of data points that has the maximum slope 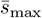 between them. The origin is taken as the midpoint between these two points as shown in Fig.(2b). This way the fitting procedure can be automated for multiple data but with reduced precision. In Fig. (2b), the fitting of *a* − *m* model (shown as *S*_*a*−*m*_) is not good and the estimated maximum growth rate is given by *m* = 0.72. This is very far from the highest slope value which is 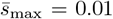. The estimated value of *m* is way different from this slope value 0.01 while using the automated fitting procedure. Whereas in Fig. (2a), with the manually selected data points, a more precise value for *m* = 0.0084 is obtained.

**Figure 1.**
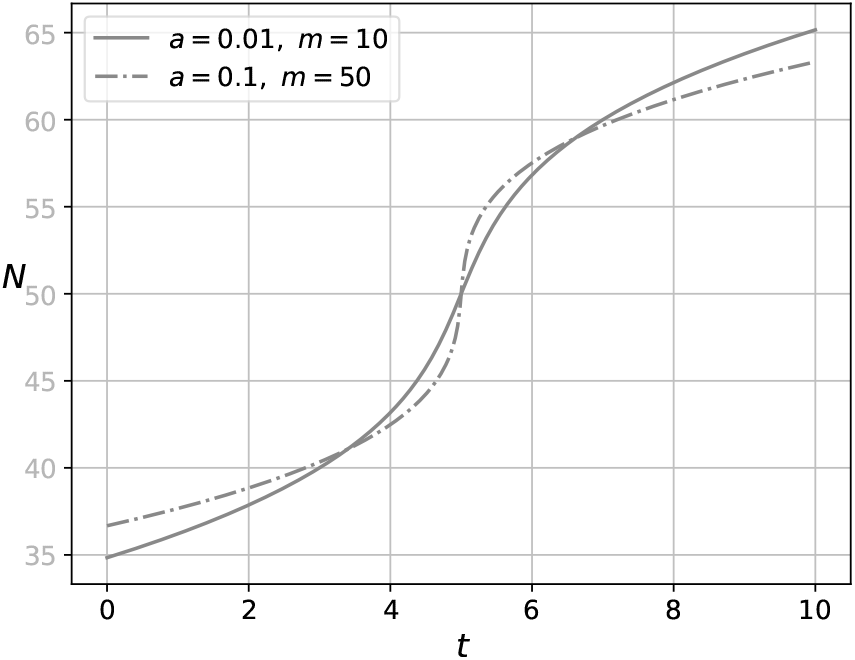
Growth curves of *a* − *m* model for various *a* and *m* values with *N*_*m*_ = 50 and *t*_*m*_ = 5.

**Figure 2.**
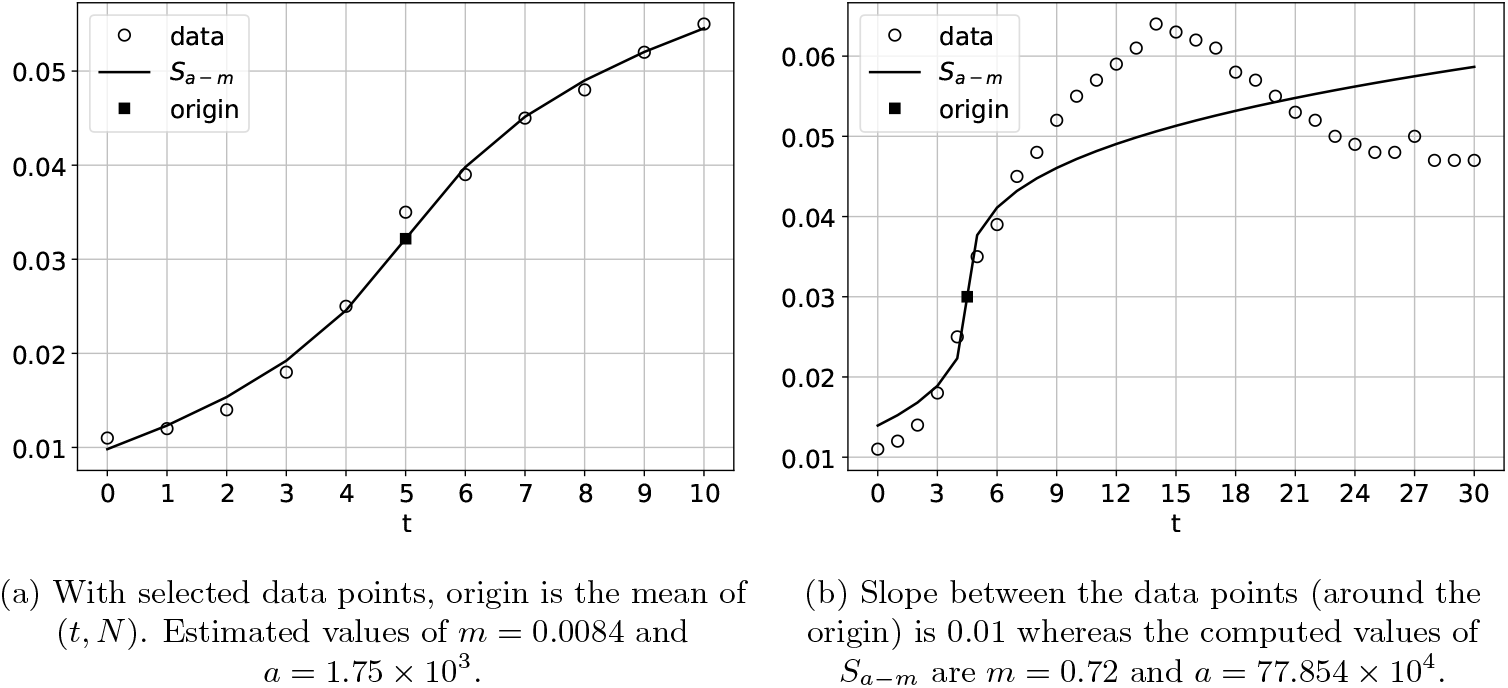
Growth curves for antibiotic concentration 0.0, shown for for strain ‘R’, replicate ‘1’.

In the following section, we introduce a procedure to get precise estimates while automating the fitting process.

## 3 Growth and decline models

The bacterial growth data includes death or decline phase [11]. So far, both the phases have been treated using separate models and the models for death phase are known as inactivation models [15]. In this work, we fit the *a* − *m* model for both of these phases.

Although the *a − m* model is a monotonically increasing function, we can use it to model the declining phase of bacterial colonies with negative values for *m* as shown in Fig. (3a). Fig. (3b) shows that it is possible to fit the death phase of bacterial growth using the *a − m* model with negative values for *m*. Based on the fit in Fig. (3b), we propose that the actual growth is a combination of growth and decline models.

**Figure 3.**
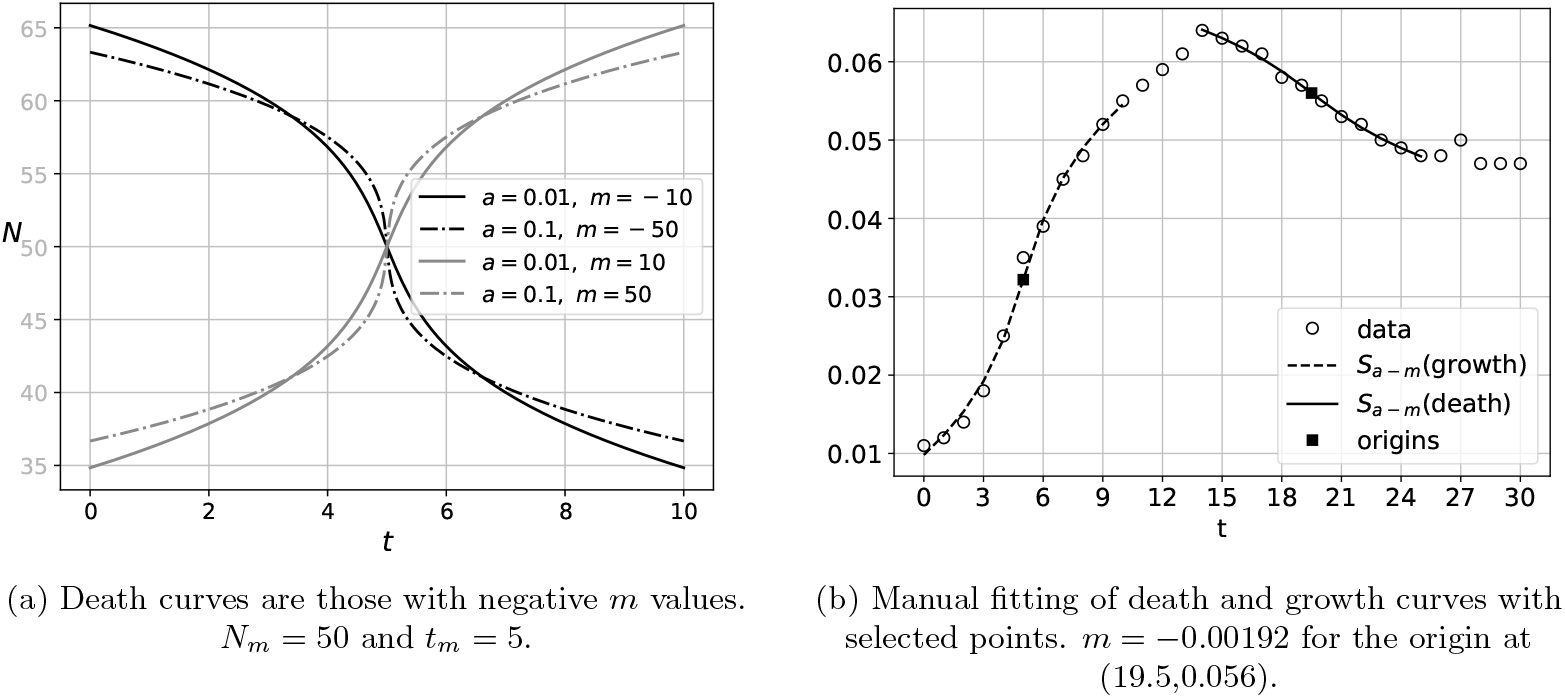
Death and growth curves of *a* − *m* model.

### 3.1 Superposition of *a* − *m* models

In order to obtain a generalized model, we represent bacterial growth to be a weighted sum i.e. superposition of growing or declining *a* − *m* models (or *S*_*a*−*m*_ curves). Although superposition is valid only for linear models, we still apply the superposition principle because the *a* − *m* model is fundamentally linear and the parameter *a* accounts for the nonlinear corrections due to superposition. This assumption is further discussed in Sec. (5.1.1).

A weighted sum of *a*−*m* models of different origins (*t*_*m*1_, *N*_*m*1_) and (*t*_*m*2_, *N*_*m*2_) with corresponding growth rates *m*_1_ and *m*_2_ is expressed as

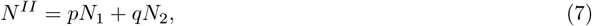

where *N*_1_ = *N* (*a, m*_1_, *t*_*m*1_, *N*_*m*1_) and *N*_2_ = *N* (*a, m*_2_, *t*_*m*2_, *N*_*m*2_) as computed from the 2^*nd*^ form (Eqn. 4) with parameters *a, m*_1_ and *a, m*_2_, respectively. Here, *p* and *q* are the weights. *a* is kept constant across the models of different origins. This is because according to the *a* − *m* model, the role of *a* is negligible near origins compared to *m*_1_ or *m*_2_. So, including *p* and *q, N*^*II*^ is a 5-parametric model. Similarly, when three *a* − *m* models are superposed as *N*^*III*^ then we will have 7 parameters (*a, m*_1_, *m*_2_, *m*_3_, *p, q, r*) and *N*^*IV*^ will be a 9-parametric model with *m*_1_, *m*_2_, *m*_3_, *m*_4_ as the growth or decline rates with *p, q, r, s* as weights. These models are fit in Fig. (4a) The growth rate of *N*_*II*_ using Eqn. (3) is given by

**Figure 4.**
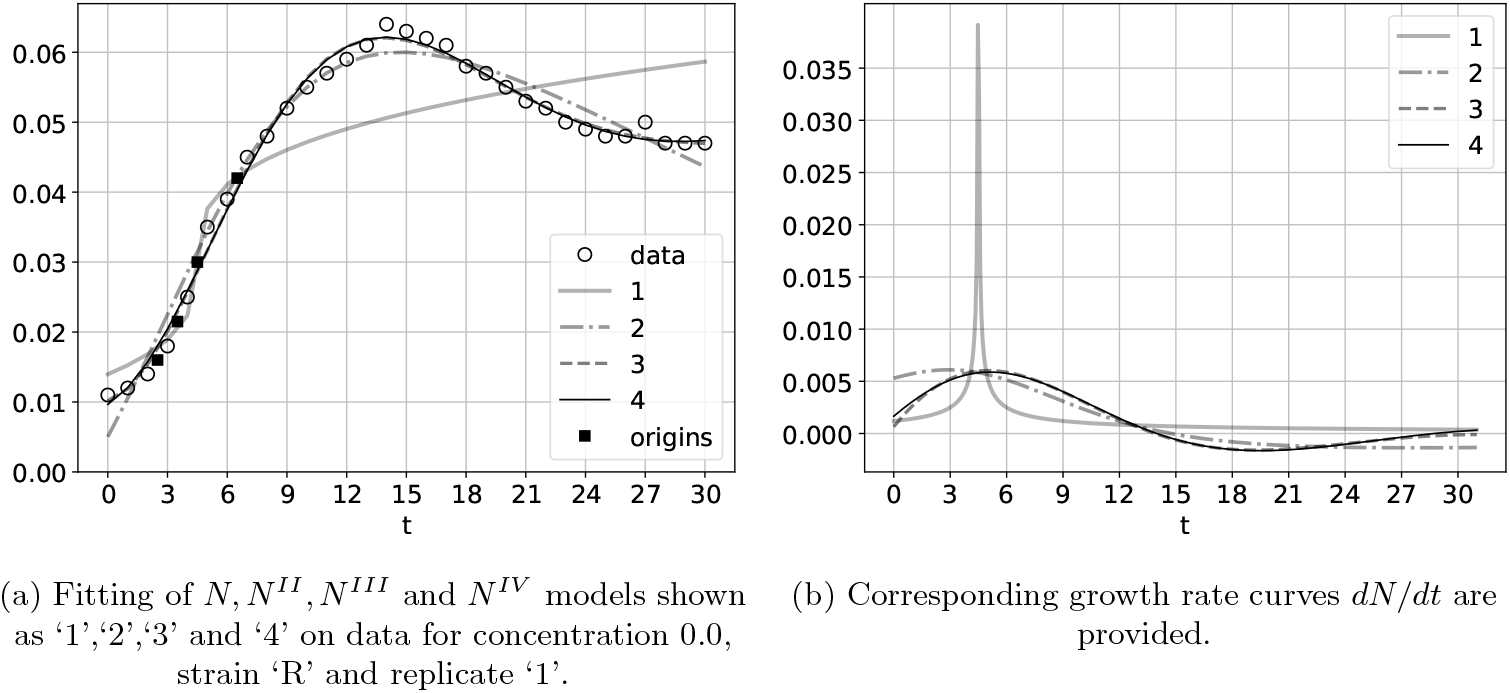
(a) Fitting of sums of *a* − *m* models with different origins and (b) the corresponding growth rates of models are shown.

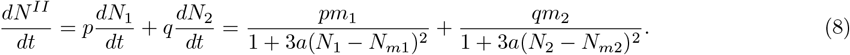

In the above equation, *N*_1_ and *N*_2_ are the values of *N* (from Eqn. (4)) for different origins with different growth or decline rates. *dN*^*III*^*/dt* and *dN*^*IV*^ */dt* can also be obtained in this way. The growth rates are compared in Fig. (4b). As the number of *a* − *m* models being superposed increases, the different curves for growth and the growth rate converge. This is evident from Fig. (4a) and Fig. (4b) where the curves of *N*^*III*^ and *N*^*IV*^ and those of *dN*^*III*^*/dt* and *dN*^*IV*^ */dt* are almost indistinguishable.

Although, the maximum growth rate *m* is different for the models in Fig. (4b), the point of maximum growth *t*_*m*_ is almost the same.

~~~
The Python implementation of the above superpostion is given by
  def N(x,a,m):
  haty = (-27*x*m/2/a+np.sqrt((27*x*m/2/a)**2+27/a**3))
return haty**(-1./3)/a-haty**(1./3)/3
#(xc1,yc1),(xc2,yc2),(xc3,yc3),(xc4,yc4) are origins
def NI(x,a,m1,xc1,yc1):
  return N(x-xc1,a,m1)+yc1
def NII(x,a,m1,m2,p,q,xc1,yc1,xc2,yc2):
 return p*(N(x-xc1,a,m1)+yc1)+q*(N(x-xc2,a,m2)+yc2)
def NIII(x,a,m1,m2,m3,p,q,r,xc1,yc1,xc3,yc2,xc3,yc3):
  return p*(N(x-xc1,a,m1)+yc1)+q*(N(x-xc2,a,m2)+yc2)+r*(
N(x-xc3,a,m3)+yc3)
def NIV(x,a,m1,m2,m3,m4,p,q,r,s,xc1,yc1,xc3,yc2,xc3,yc3,xc4,yc4):
  return p*(N(x-xc1,a,m1)+yc1)+q*(N(x-xc2,a,m2)+yc2)+r*( N(x-xc3,a,m3)+yc3)+s*(N(x-xc4,a,m4)+yc4)
~~~

The derivatives are implemented as:

~~~
def Ngrad(x,a,m):
  return m/(1+3*a*N(x,a,m)**2)
NIIgrad = p*Ngrad(xder-xc1,a,m1) + q*Ngrad(x-xc2,a,m2)
NIIIgrad = p*Ngrad(x-xc1,a,m1) + q*Ngrad(x-xc2,a,m2) + r*Ngrad(x-xc3,a,m3)
NIVgrad = p*Ngrad(x-xc1,a,m1) + q*Ngrad(x-xc2,a,m2) + r*Ngrad(x-xc3,a,m3) + s*Ngrad(x-xc4,a,m4)
~~~

## 4 Results and discussions

Given below are plots of bacterial growth and growth rates for different antibiotic concentrations. The initial guesses are

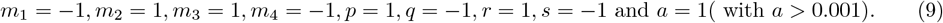

1. Fig. (5) for antibiotic concentration 0: All the three strains show S-curve growth pattern. For ‘R1’ strain, the decline phase too is traced by the curves of *NII, NIII* and *NIV*. Also, the initial peak of growth rate is well-captured by the growth rate curves given at the bottom of each growth plot. *N*^*IV*^ captures the initial lag phase better than the rest of the models.
2. Fig. (6) for concentration 0.24: At this concentration, the distinct growth rate peaks shown by ‘R1’ and ‘R2’ for zero concentration disappear. Although, *N*^*IV*^ captures the lag phase for ‘T1’, ‘T2’ and ‘D2’, it is the single origin (N) *a* − *m* model that captures the lag phase for ‘D1’.
3. Fig. (7) for concentration 0.49: Although *N*^*IV*^ captures the data well, *N*^*III*^ peaks at the expected time for ‘D1’ and ‘D2’.
4. Fig. (8) for concentration 0.98: ‘T2’ has a small but distinct decline phase which is captured by *N*^*III*^ and *N*^*IV*^ growth rate curves. For ‘D1’ and ‘D2’, only *N*^*IV*^ captures the expected growth rate peak.
5. Fig. (9) for concentration 1.95: For the ‘T2’ data, only *N*^*III*^ captures the maximum growth rate well than the rest of the models. However, *N*^*IV*^ captures well for the rest of the data.
6. Fig. (10) for concentration 3.91: *N*^*IV*^ captures the lag phase better than the rest of the models.
7. Fig. (11) for concentration 7.81: Only *N*^*IV*^ captures the peaking of growth rate for ‘D1’.
8. Fig. (12) for concentration 15.63: *N*^*III*^ captures the lag phase and the peaking of growth rate for ‘T2’, whereas, *N*^*IV*^ fits better for ‘D2’
9. Fig. (13) for concentration 31.25: Overall, *N*^*III*^ performs better than *N*^*IV*^. *N*^*IV*^ overfits the data for ‘D1’ showing a declining peak in the growing phase.
10. Fig. (14) for concentration 62.5:While *N*^*IV*^ fits well for ‘T2’, *N*^*III*^ fits better for ‘T1’. The maximum growth rate peak is however, captured by *N*^*IV*^ for ‘T1’ at *t <* 3.
11. Fig. (15) for concentration 125: *N*^*II*^ and *N*^*III*^ fit better than *N*^*IV*^ for ‘R1’.
12. Fig. (16) for concentration 250: At this concentration, all the three strains show a slump in growth. But surprisingly a peak is captured for ‘R1’ by the *N*^*IV*^ model. Other than this ‘R1’ peaks only at concentration 0 and 31.5.

**Figure 5.**
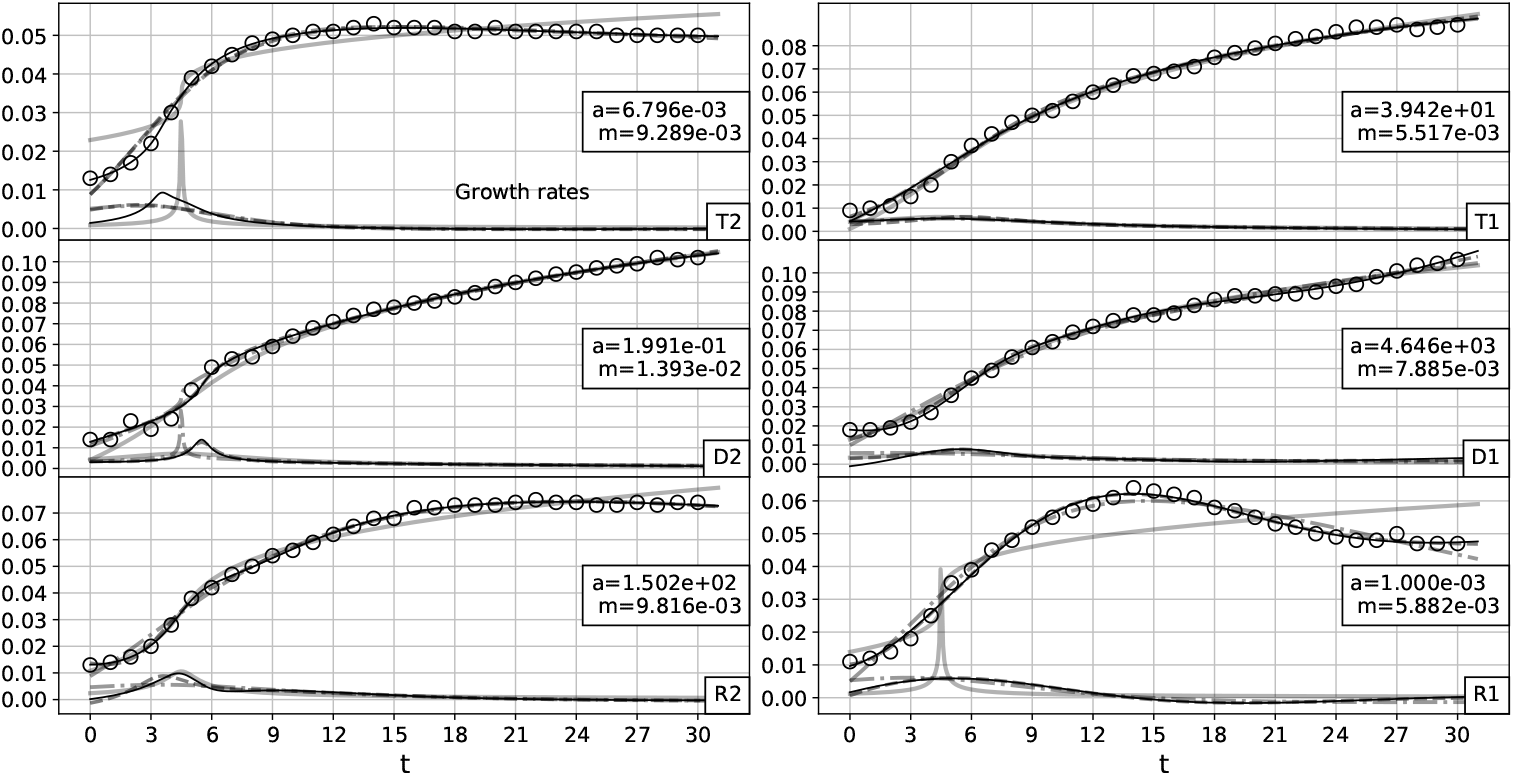
Growth and growth rate curves for antibiotic concentration 0.0, shown for three strains ‘T’, ‘D’ and ‘R’ and for two replicates ‘1’ and ‘2’ as ‘T1’,’T2’,’D1’,’D2’,’R1’ and ‘R2’. The growth rate curves are shown at the bottom of the growth curves. *a* and *m* values are provided for *N*^*IV*^. *N, N*^*II*^, *N*^*III*^ and *N*^*IV*^ are also shown using the same legends of Fig. (4a) and (4b).

**Figure 6.**
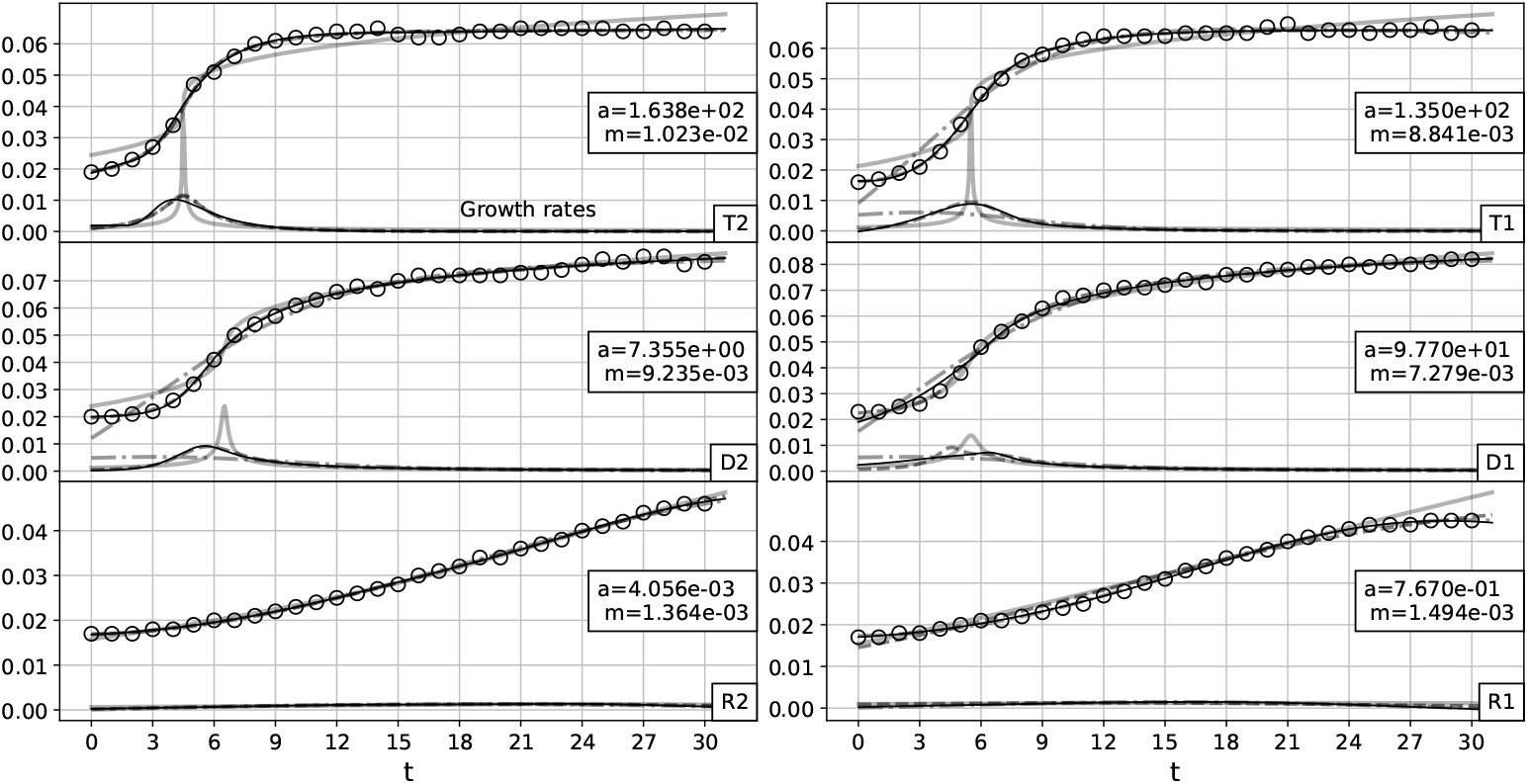
Growth and growth rate curves for antibiotic concentration 0.24, shown for three strains ‘T’, ‘D’ and ‘R’ and for two replicates ‘1’ and ‘2’ as ‘T1’,’T2’,’D1’,’D2’,’R1’ and ‘R2’. The growth rate curves are shown at the bottom of the growth curves. *a* and *m* values are provided for *N*^*IV*^. *N, N*^*II*^, *N*^*III*^ and *N*^*IV*^ are also shown using the same legends of Fig. (4a) and (4b).

**Figure 7.**
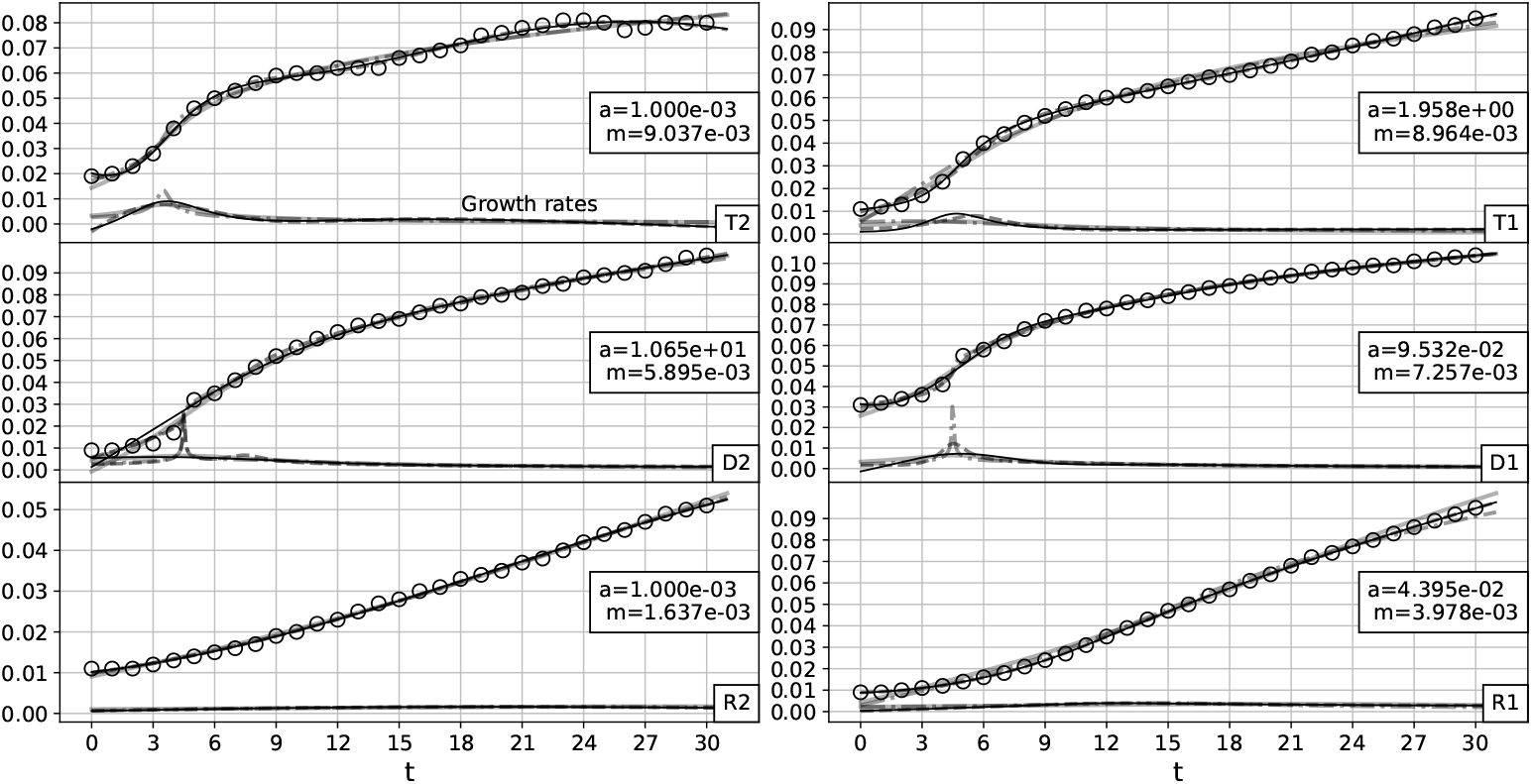
Growth and growth rate curves for antibiotic concentration 0.49, shown for three strains ‘T’, ‘D’ and ‘R’ and for two replicates ‘1’ and ‘2’ as ‘T1’,’T2’,’D1’,’D2’,’R1’ and ‘R2’. The growth rate curves are shown at the bottom of the growth curves. *a* and *m* values are provided for *N*^*IV*^. *N, N*^*II*^, *N*^*III*^ and *N*^*IV*^ are also shown using the same legends of Fig. (4a) and (4b).

**Figure 8.**
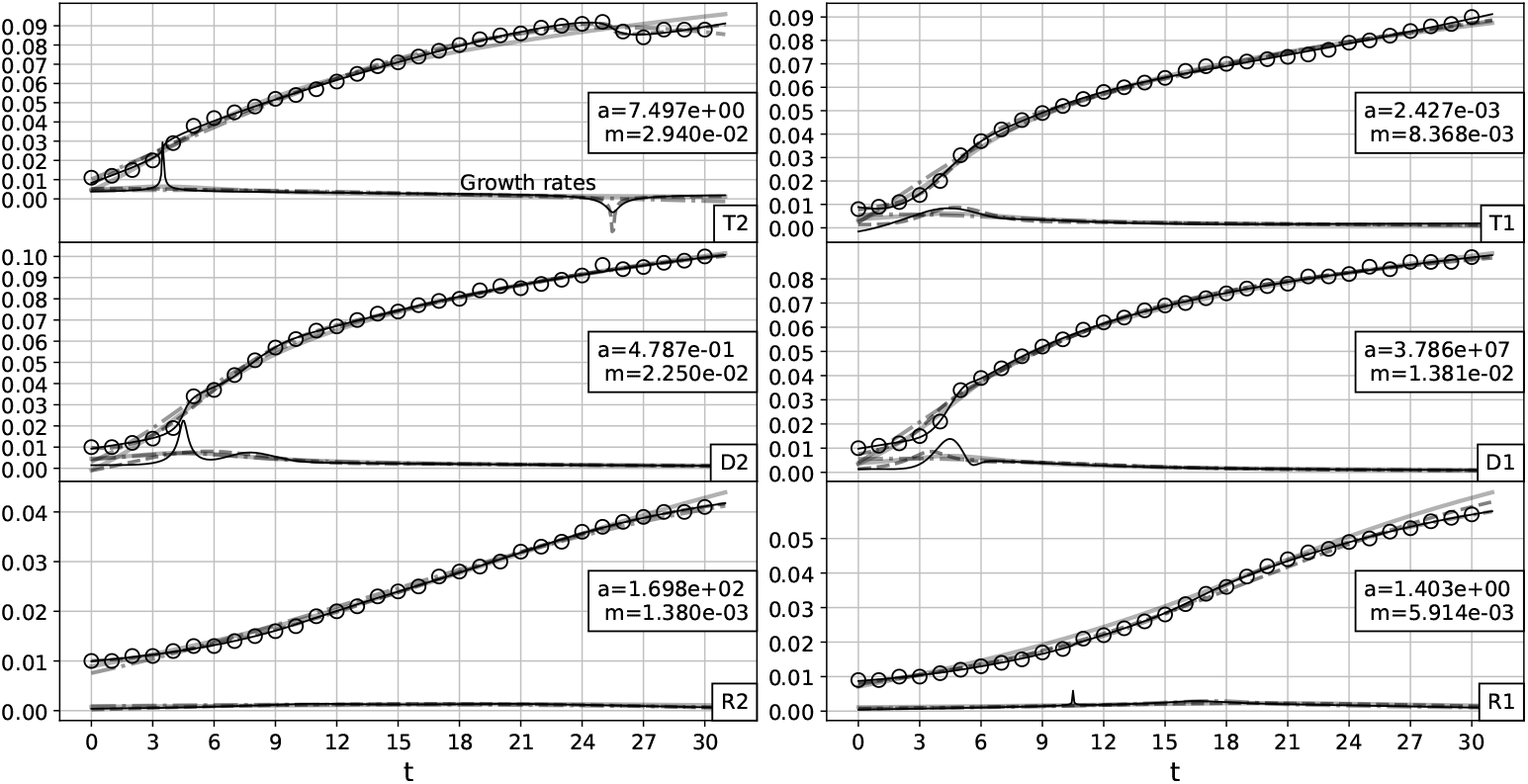
Growth and growth rate curves for antibiotic concentration 0.98, shown for three strains ‘T’, ‘D’ and ‘R’ and for two replicates ‘1’ and ‘2’ as ‘T1’,’T2’,’D1’,’D2’,’R1’ and ‘R2’. The growth rate curves are shown at the bottom of the growth curves. *a* and *m* values are provided for *N*^*IV*^. *N, N*^*II*^, *N*^*III*^ and *N*^*IV*^ are also shown using the same legends of Fig. (4a) and (4b).

**Figure 9.**
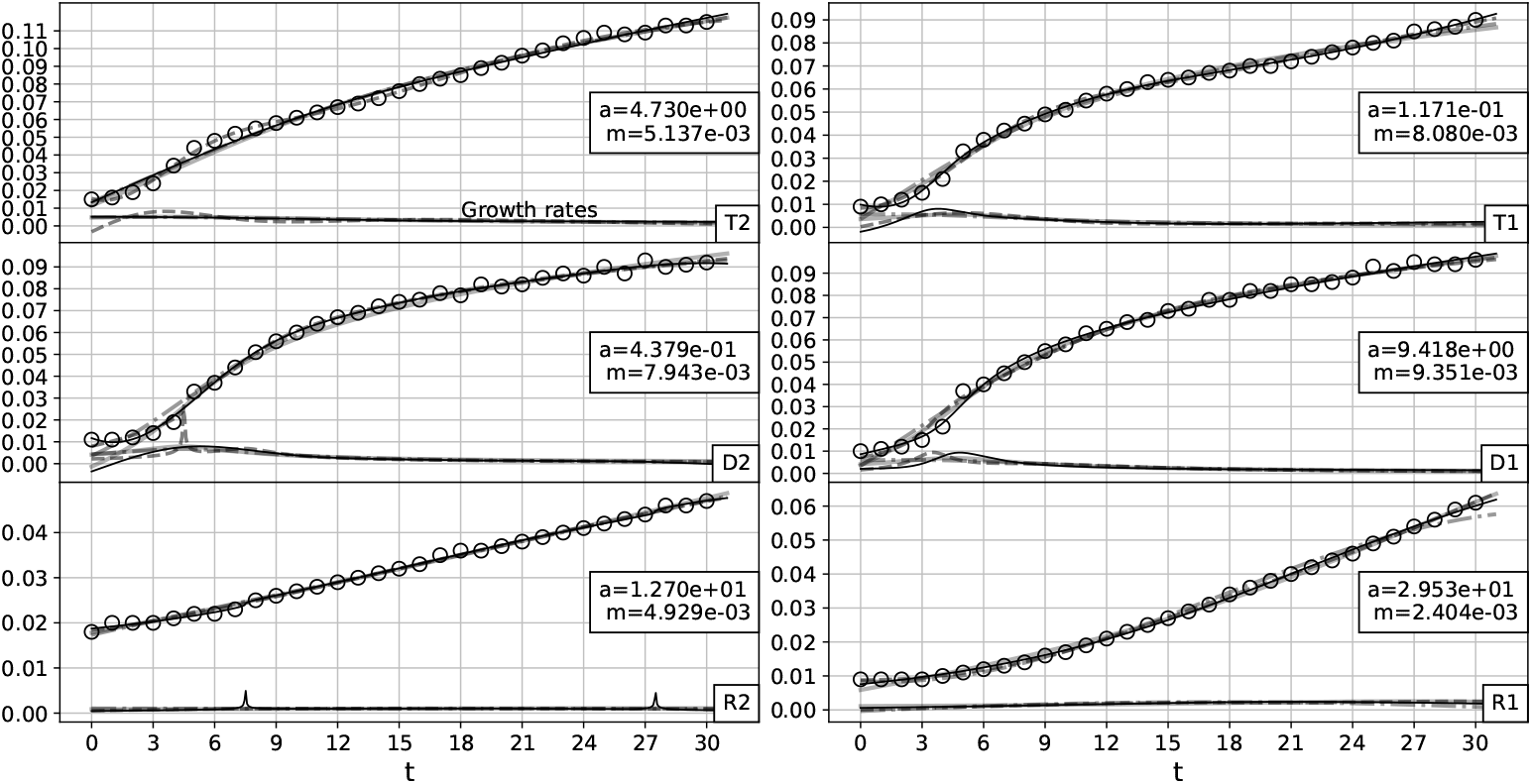
Growth and growth rate curves for antibiotic concentration 1.95, shown for three strains ‘T’, ‘D’ and ‘R’ and for two replicates ‘1’ and ‘2’ as ‘T1’,’T2’,’D1’,’D2’,’R1’ and ‘R2’. The growth rate curves are shown at the bottom of the growth curves. *a* and *m* values are provided for *N*^*IV*^. *N, N*^*II*^, *N*^*III*^ and *N*^*IV*^ are also shown using the same legends of Fig. (4a) and (4b).

**Figure 10.**
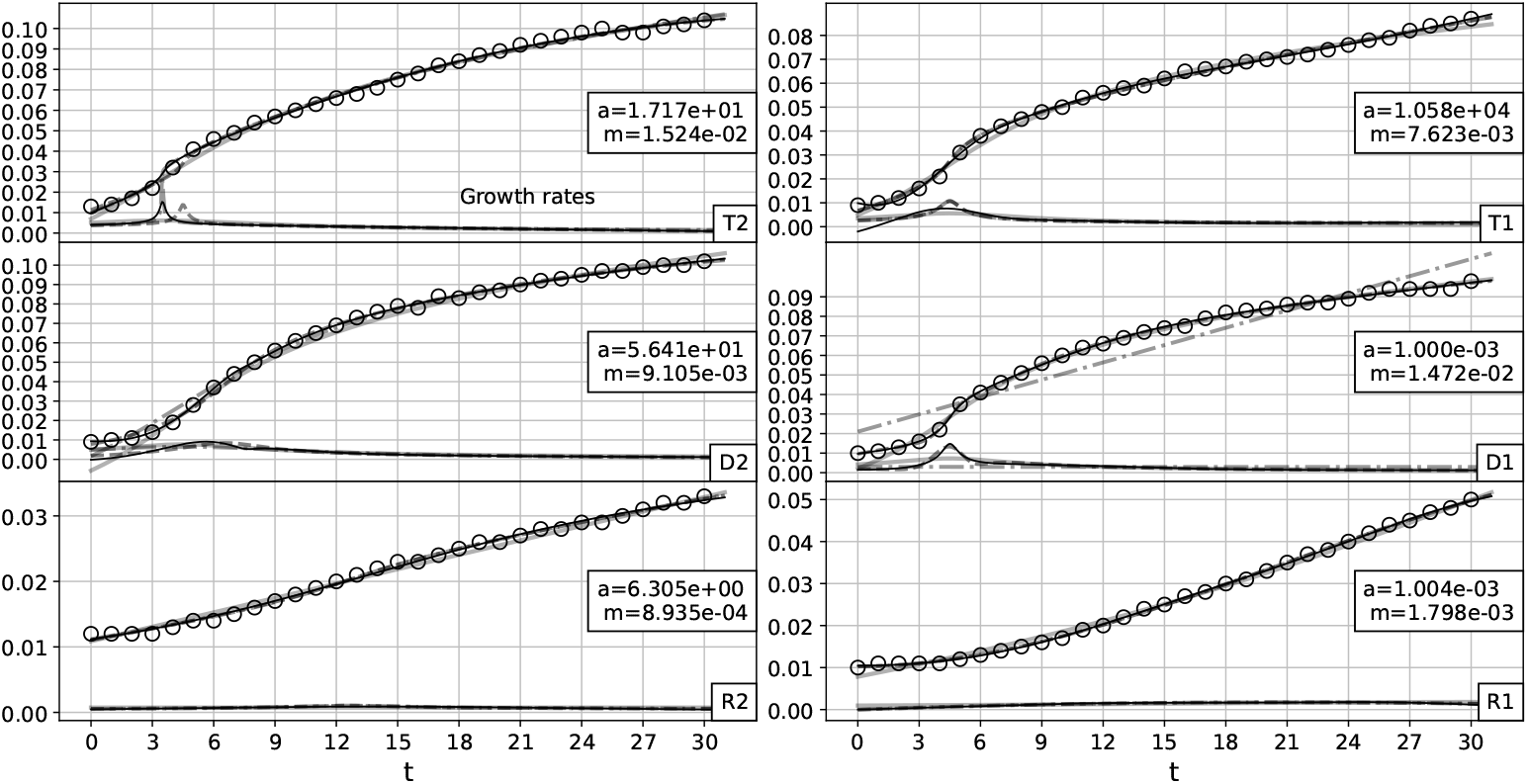
Growth and growth rate curves for antibiotic concentration 3.91, shown for three strains ‘T’, ‘D’ and ‘R’ and for two replicates ‘1’ and ‘2’ as ‘T1’,’T2’,’D1’,’D2’,’R1’ and ‘R2’. The growth rate curves are shown at the bottom of the growth curves. *a* and *m* values are provided for *N*^*IV*^. *N, N*^*II*^, *N*^*III*^ and *N*^*IV*^ are also shown using the same legends of Fig. (4a) and (4b).

**Figure 11.**
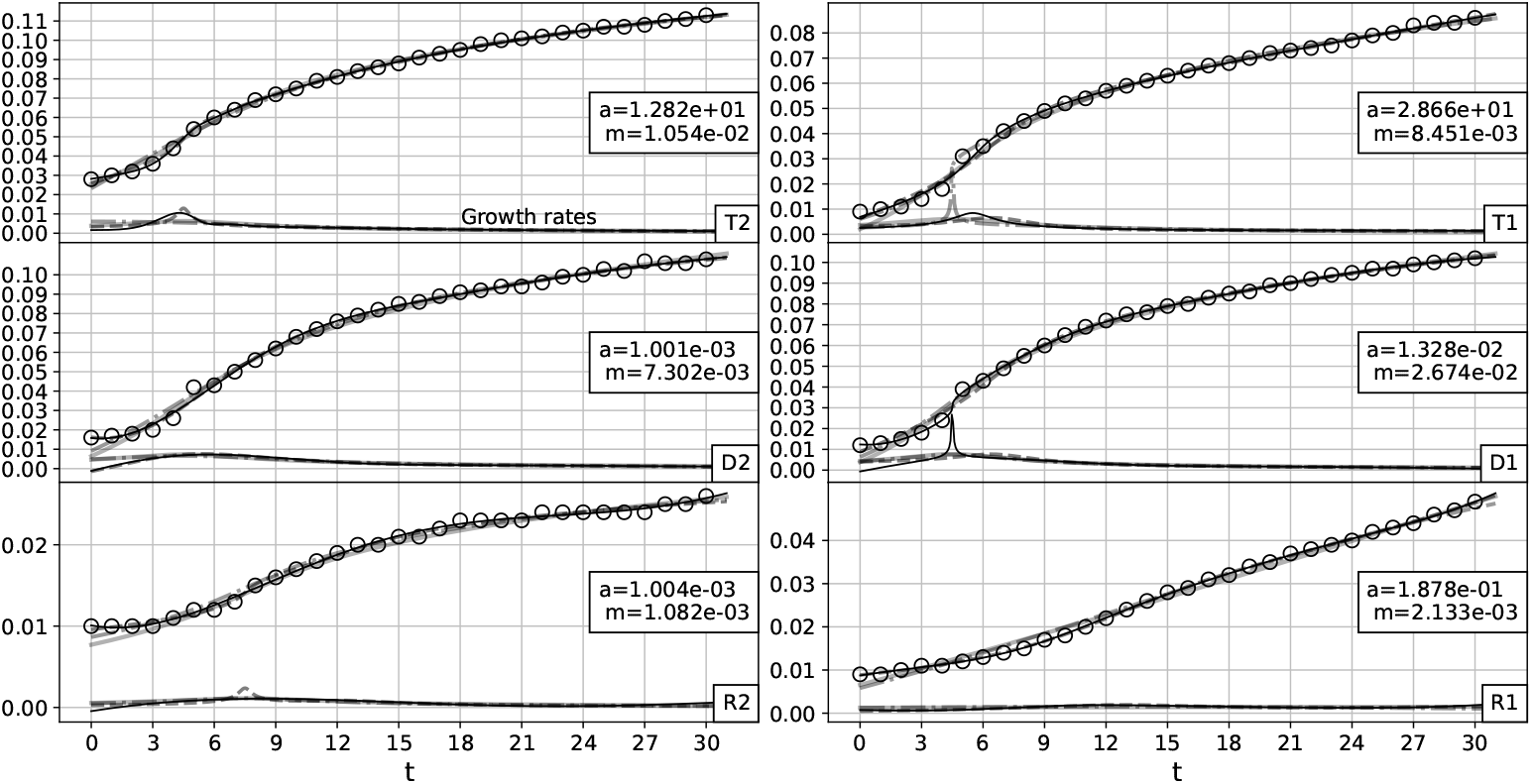
Growth and growth rate curves for antibiotic concentration 7.81, shown for three strains ‘T’, ‘D’ and ‘R’ and for two replicates ‘1’ and ‘2’ as ‘T1’,’T2’,’D1’,’D2’,’R1’ and ‘R2’. The growth rate curves are shown at the bottom of the growth curves. *a* and *m* values are provided for *N*^*IV*^. *N, N*^*II*^, *N*^*III*^ and *N*^*IV*^ are also shown using the same legends of Fig. (4a) and (4b).

**Figure 12.**
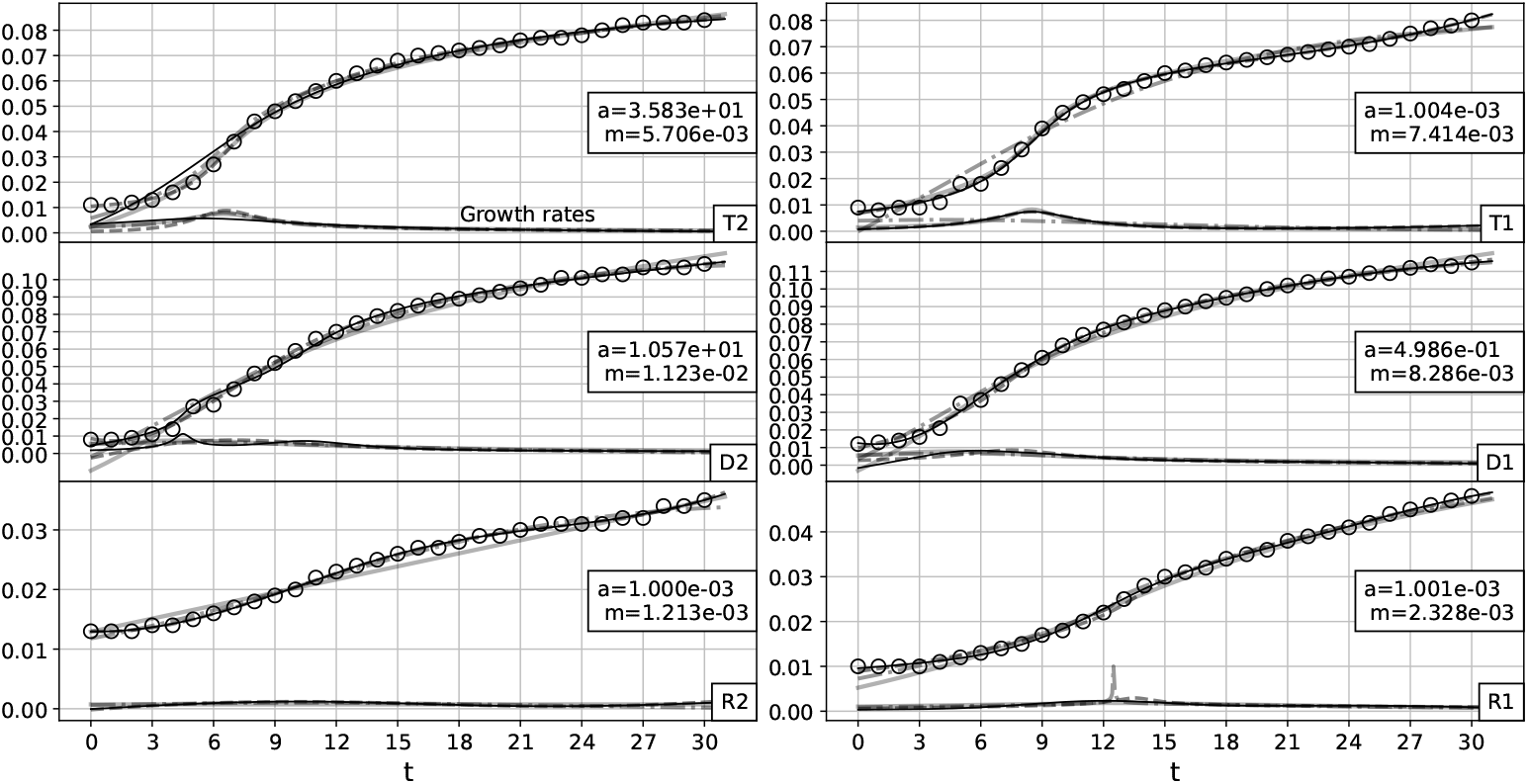
Growth and growth rate curves for antibiotic concentration 15.63, shown for three strains ‘T’, ‘D’ and ‘R’ and for two replicates ‘1’ and ‘2’ as ‘T1’,’T2’,’D1’,’D2’,’R1’ and ‘R2’. The growth rate curves are shown at the bottom of the growth curves. *a* and *m* values are provided for *N*^*IV*^. *N, N*^*II*^, *N*^*III*^ and *N*^*IV*^ are also shown using the same legends of Fig. (4a) and (4b).

**Figure 13.**
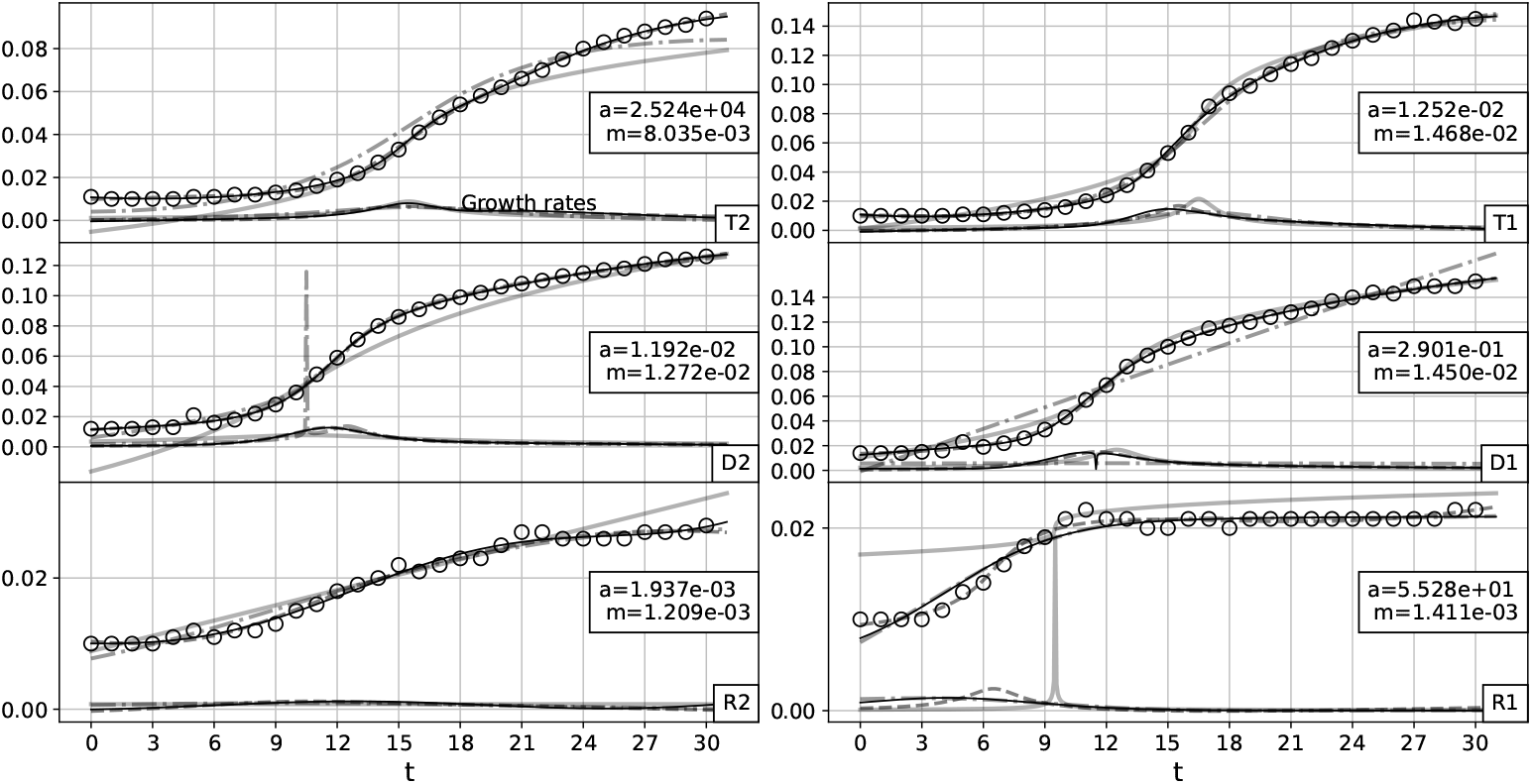
Growth and growth rate curves for antibiotic concentration 31.25, shown for three strains ‘T’, ‘D’ and ‘R’ and for two replicates ‘1’ and ‘2’ as ‘T1’,’T2’,’D1’,’D2’,’R1’ and ‘R2’. The growth rate curves are shown at the bottom of the growth curves. *a* and *m* values are provided for *N*^*IV*^. *N, N*^*II*^, *N*^*III*^ and *N*^*IV*^ are also shown using the same legends of Fig. (4a) and (4b).

**Figure 14.**
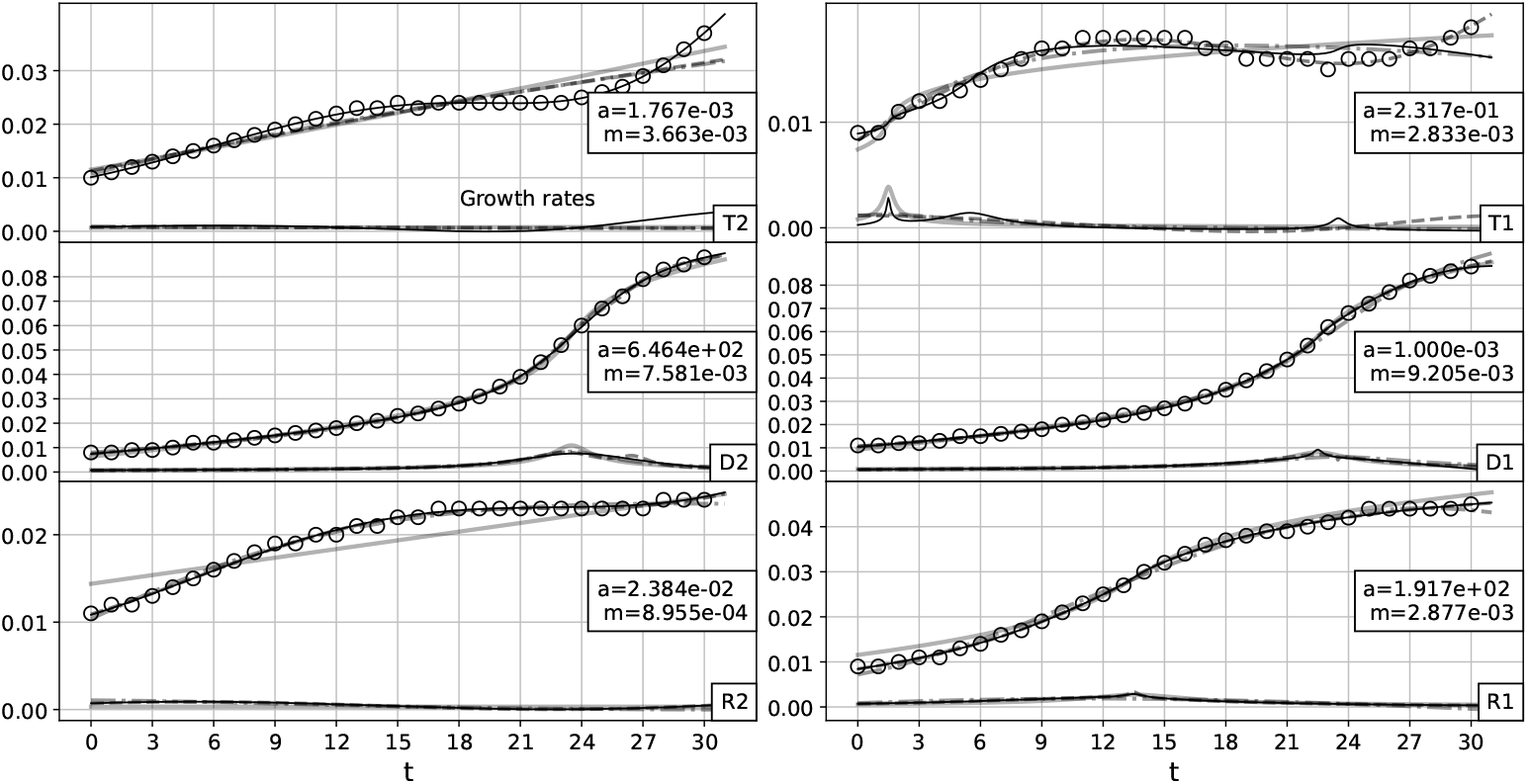
Growth and growth rate curves for antibiotic concentration 62.5, shown for three strains ‘T’, ‘D’ and ‘R’ and for two replicates ‘1’ and ‘2’ as ‘T1’,’T2’,’D1’,’D2’,’R1’ and ‘R2’. The growth rate curves are shown at the bottom of the growth curves. *a* and *m* values are provided for *N*^*IV*^. *N, N*^*II*^, *N*^*III*^ and *N*^*IV*^ are also shown using the same legends of Fig. (4a) and (4b).

**Figure 15.**
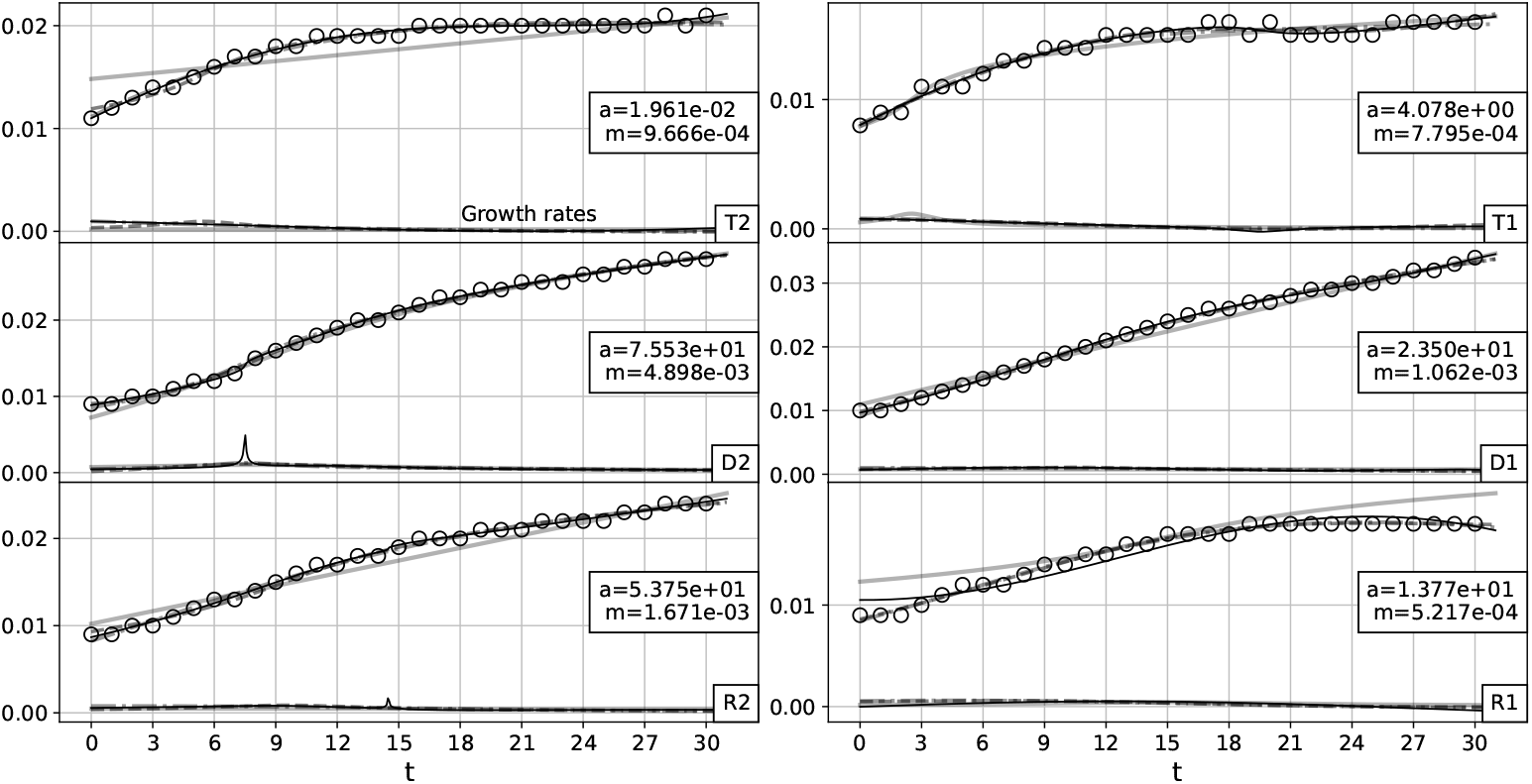
Growth and growth rate curves for antibiotic concentration 125.0, shown for three strains ‘T’, ‘D’ and ‘R’ and for two replicates ‘1’ and ‘2’ as ‘T1’,’T2’,’D1’,’D2’,’R1’ and ‘R2’. The growth rate curves are shown at the bottom of the growth curves. *a* and *m* values are provided for *N*^*IV*^. *N, N*^*II*^, *N*^*III*^ and *N*^*IV*^ are also shown using the same legends of Fig. (4a) and (4b).

**Figure 16.**
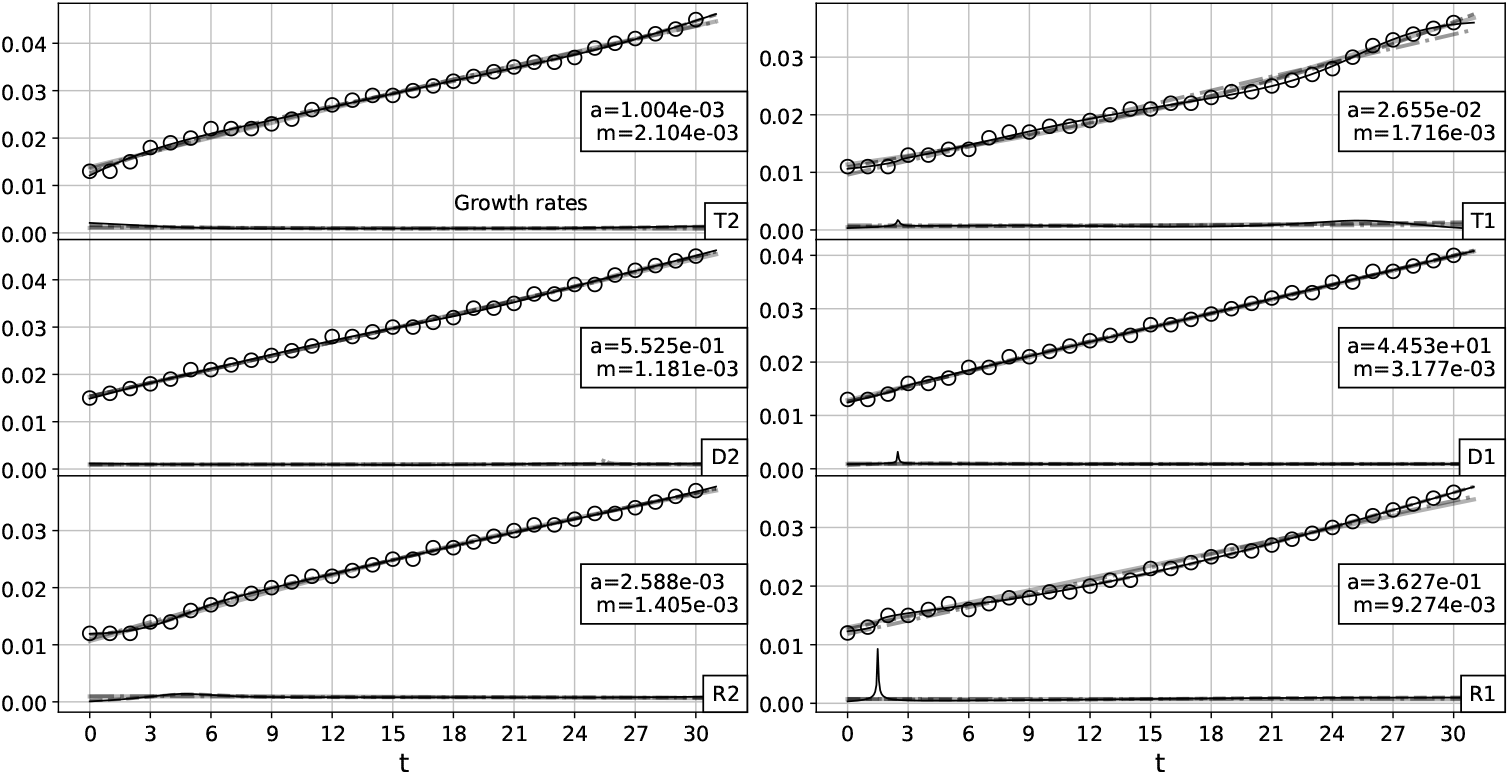
Growth and growth rate curves for antibiotic concentration 250.0, shown for three strains ‘T’, ‘D’ and ‘R’ and for two replicates ‘1’ and ‘2’ as ‘T1’,’T2’,’D1’,’D2’,’R1’ and ‘R2’. The growth rate curves are shown at the bottom of the growth curves. *a* and *m* values are provided for *N*^*IV*^. *N, N*^*II*^, *N*^*III*^ and *N*^*IV*^ are also shown using the same legends of Fig. (4a) and (4b).

### 4.1 A growth coordinate system

The values of *a* and *m* for models *N*^*III*^ and *N*^*IV*^ obtained from Figs. (6) to (16) are compared on *xy*− plane in Figs. (17) and (18) with log *a* as the *x* − *axis* and *m* as the *y*− axis [7]. Thus, the entire data of plate reader experiments can be conveniently compared using the *a* − *m* growth coordinate system. Some of the observations from the coordinate system plots are:

**Figure 17.**
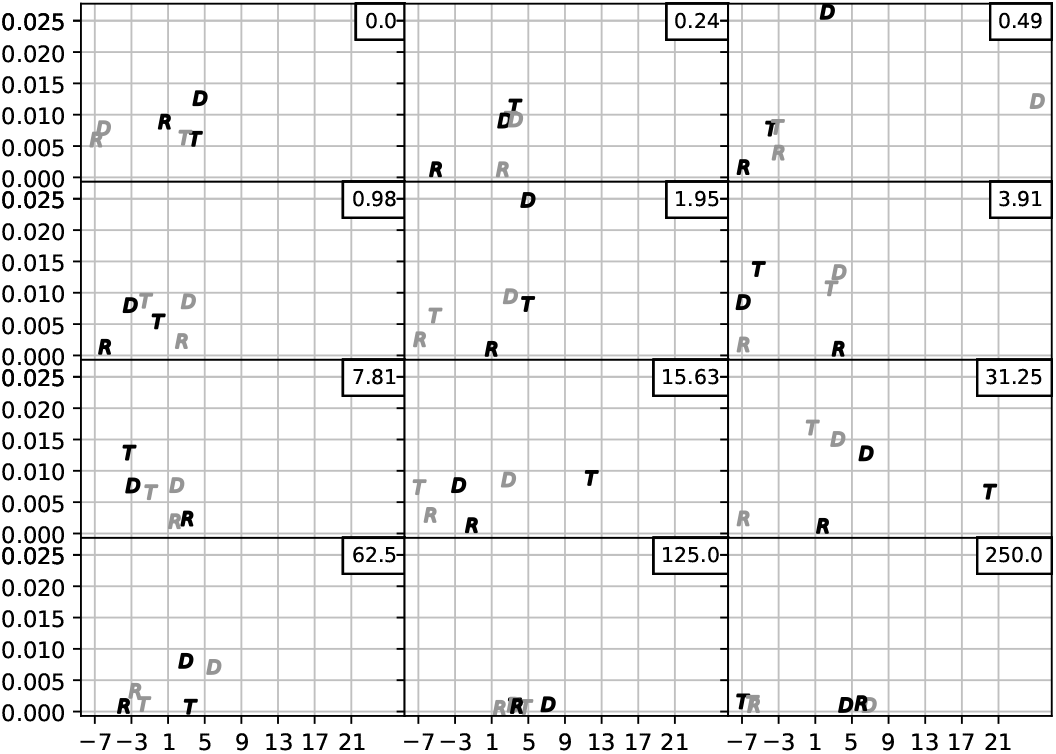
Comparison of bacterial growth using a growth coordinate system with log *a* as the *x* − axis and *m* as the *y* − axis. *a* and *m* values are obtained using *N*^*III*^ model. The grey markers are (log *a, m*) of replicate ‘1’ and the black markers are those of replicate ‘2’. Concentration values are shown as numerical values within the frame at the top right side of each plot.

**Figure 18.**
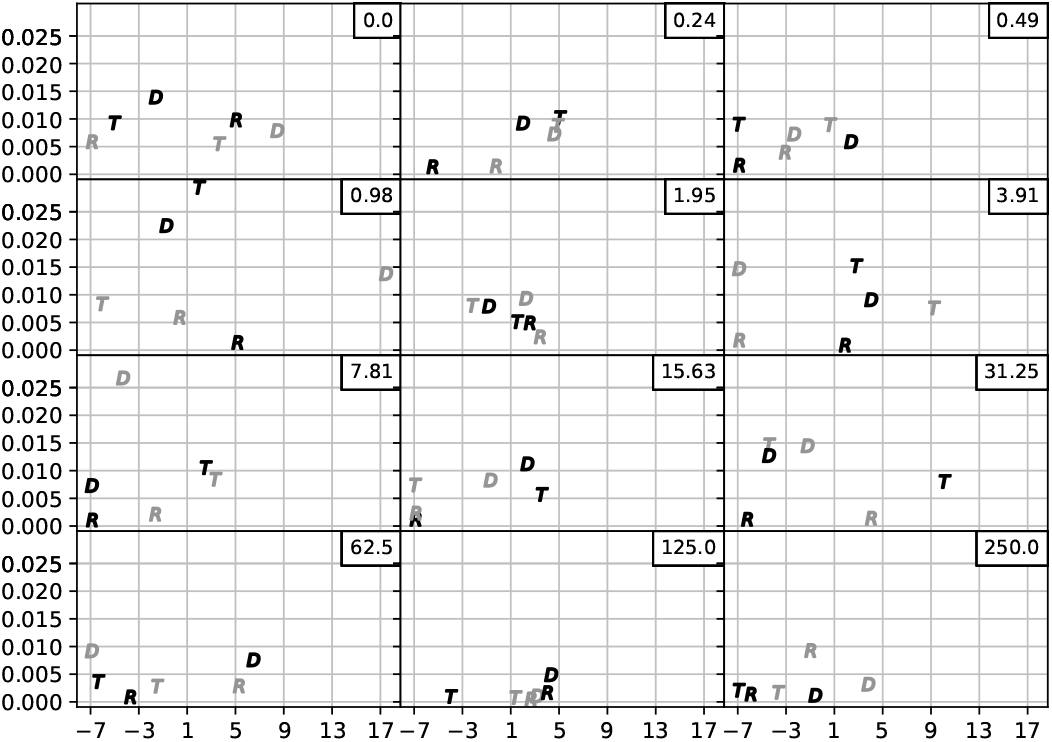
Comparison of bacterial growth using a growth coordinate system with log *a* as the *x* − axis and *m* as the *y* − axis. *a* and *m* values are obtained using *N*^*IV*^ model. The grey markers are (log *a, m*) of replicate ‘1’ and the black markers are those of replicate ‘2’. Concentration values are shown as numerical values within the frame at the top right side of each plot.

1. At concentration 0.49: For the *N*^*III*^ model (Fig. (17)) the *m* value of ‘D2’ is higher than the estimate of *N*^*IV*^ model (Fig. (18)). This reflects the precise capture of *m* by the *N*^*III*^ model (as seen in Fig. (7)). ‘D1’ has a high value of *a* for the *N*^*III*^ model. It can be inferred that this is due to the increase in restrictions after the maximum growth.
2. At concentration 0.98: *N*^*IV*^ provides better estimate for *m* as seen in Fig. (8) for ‘T2’ and ‘D2’. This is reflected in Fig. (18). High *a* value for ‘D1’ is observed for the *N*^*IV*^ model which reflects high restrictions after maximum growth as observed in Fig. (8).
3. At concentration 250: *N*^*IV*^ model captures increase in *m* for ‘R1’. This can also be seen in Fig. (16).

## 5 Advantages and limitations

The *a* − *m* model has the following advantages:

1. Rather than using a set of differential equations to model growth, *a* − *m* model uses algebraic expressions. Thus, differential equation solver packages are avoided to fit the data.
2. Both the growth and decline phase are modeled together with the *a* − *m* model.
3. Although more than two parameters are used to fit the data, the models *N*^*II*^, *N*^*III*^ and *N*^*IV*^ can be interpreted using a coordinate system with just two parameters *a* and *m*.
4. Growth rates are directly obtained from the models. Thus, the distribution of growth rates over time is also provided in Figs. (5) - (16).
5. Moreover, the variance due to nonlinearity is accounted for with just one parameter *a*. Whereas, when using higher order polynomials in *x*, the model may overfit the data [2]. The overfitting problem is avoided to some extent with *a* − *m* models.

### 5.1 Limitations

#### 5.1.1 Validating superposition

The net linear maximum growth rate for an *N*^*II*^ model would be

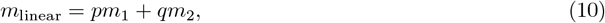

which is independent of *a*, i.e., independent of nonlinear restrictions. The value of above linear sum is compared with the observed maximum value of *dN*^*II*^*/dt* (Eqn. (8)) which includes nonlinear terms through *a*. Therefore, the difference

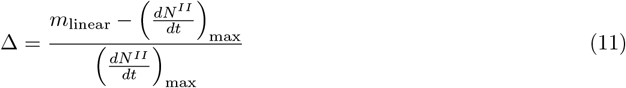

indicates the influence of nonlinearities. This relative difference is less than 5% for 60% of the *N*^*II*^ models, 51% of the *N*^*III*^ models and 25% of the *N*^*IV*^ models. As we superpose more *a* − *m* models the estimated *m* deviates away from *m*_linear_ due to its dependence on *a*. This deviation may also depend on initial conditions Eqn. (9) that was set while fitting the models. Thus, modeling based on several initial guesses may be required to understand and characterize the deviation due to nonlinearity.

#### 5.1.2 Dependence on initial conditions

The models *N*^*II*^, *N*^*III*^ and *N*^*IV*^ need specific initial conditions. *N*^*II*^ model does not fit the data in ‘D1’ of Figs. (10) and (13) compared to *N* which is the single origin *a* − *m* model. This is because the initial guess (Eqn. (9)) is not appropriate for that particular data. However, if the initial guess is changed the *N*^*II*^ model fits better than *N*.

## 6 Conclusions

In this work, we model both growth and decline using continued fraction of straight lines. To fit the growth data, we use the principle of superposition and sum the *a* − *m* models of different origins while accounting for nonlinear corrections using *a*. The sums of models are applicable on large-scale growth data. Since analytically tractable derivative expressions are available, the growth rates are obtained directly after fitting the models on the data. The models and their results are also compared using a planar growth coordinate system. Thus, the nonlinear dynamics of bacterial growth can be captured, analyzed and interpreted using *a* − *m* models and their sums. Moreover, the chosen origins of models reveal the importance of certain time intervals in the growth data. Although the growth data is obtained for several hours of experiments, events at these specific time intervals seem to decide the entire growth pattern.

## Code and Supplementary material

The code and estimations for models *N, N*^*II*^, *N*^*III*^ and *N*^*IV*^ are provided at the github repository: grasshopper14/a-m-bacterial-growth

## Acknowledgement

We thank researchers such as Claudia Seiler for making the data publicly available. It is of great help especially for independent researchers like us.

